# Telocyte networks are an essential stem cell niche component in hair follicle regeneration

**DOI:** 10.1101/2023.05.17.541070

**Authors:** Marco Canella, Simcha Nalick, Amal Gharbi, Abrar Jamous, Noa Corem, Ittai Ben-Porath, Matan Hofree, Michal Shoshkes-Carmel

**Affiliations:** Department of Developmental Biology and Cancer Research, Institute for Medical Research Israel-Canada, Hebrew University Medical School, Jerusalem, Israel; The Benin School of Computer Science and Engineering, The Hebrew University of Jerusalem, Israel; Lautenberg Centre for Immunology and Cancer Research, Institute for Medical Research Israel-Canada, Hebrew University Medical School, Jerusalem, Israel

**Keywords:** stem cells, stem-cell niche, mesenchyme, telocytes, hair follicle

## Abstract

Self-renewing tissues rely on stem cells (SCs) to regenerate differentiated cells, requiring precise coordination between SCs and their progeny. Here, we investigated how SC activity is regulated across multiple epithelial layers, using the hair follicle (HF) as a model system. We uncovered an extensive, interconnected network of telocytes, specialized mesenchymal cells, that spans all HF layers, including regions previously thought to lack mesenchymal or connective tissue components. These telocyte networks remain in constant contact with SCs and their progeny throughout regeneration, dynamically adapting their structure and molecular profile to provide localized, phase-specific signals. By employing two independent mouse models to ablate telocytes or disrupt their Wnt signaling, we demonstrate that telocyte networks are an essential component of the SC niche. Our findings propose a revised, integrative model of the SC niche, in which telocyte networks and their Wnt production play a central role in adult SC biology and tissue regeneration.

## Introduction

A key question in tissue regeneration is how stem cell (SC) regulation is coordinated with the control of their progeny to ensure a balanced production of new cells and maintain homeostasis. The intestine serves as a model of continuous regeneration, where SC differentiation follows a linear process along the crypt-villus axis, forming a single-cell layer epithelium supported by connective tissue. Recent studies in the intestine have identified a subepithelial network of specialized mesenchymal cells called telocytes^1–5^, which express Foxl1 and are crucial components of the SC niche^6–9^. These cells provide Wnt proteins that drive the proliferation of stem and progenitor cells^6,7^, while also facilitating enterocyte differentiation^10^, potentially linking SC regulation with progeny differentiation and playing a significant role in maintaining tissue homeostasis^7,8,10,11^.

In contrast to the intestine, the hair follicle exhibits a more complex, nonlinear, and dynamic differentiation process within its outer and inner concentric epithelial layers^12^. The follicle undergoes cycles of rest, growth and regression, involving significant morphological and molecular changes that require precise, dynamic control.

Mice are an ideal model for studying these dynamics due to their synchronized initial hair cycles. Hair follicle SCs reside in the basal layer of the bulge, just below the sebaceous gland at the follicle’s lower permanent end. These SCs are typically slow-cycling and label-retaining^13–16^. The inner layer of the bulge, formed by K6 differentiated SC progeny, is thought to create a niche that inhibits SC function^12^. Beneath the bulge, a small group of cells known as the hair germ proliferates more actively and initiates follicle growth^12,16–21^. During the transition from the resting to the growth phase, hair germ cells are stimulated first, followed by bulge cell proliferation, driving the hair cycle forward^17^. The hair germ matures into transit-amplifying matrix cells, which rapidly proliferate at the follicle’s bulb and differentiate into inner root sheath cells, contributing to hair formation. Meanwhile, bulge SCs primarily generate relatively undifferentiated basal outer root sheath cells, which retain many stem cell characteristics. These cells accelerate proliferation as they approach the follicle bulb^12^.

Throughout most of the hair growth cycle, follicle SCs remain quiescent, activated only at the cycle’s onset. Bone morphogenetic proteins (BMPs) maintain this quiescence^12,22–24^, while the β-catenin-mediated Wnt-signaling pathway is crucial for initiating SC activation and growth^17,25–30^. The absence of β-catenin leads to follicular arrest in the resting phase^25,31^.

During the regeneration cycle, the lower portion of the hair follicle remains in close contact with a cluster of mesenchymal cells known as the dermal papilla. Since the 1960s, studies on vibrissa follicles have demonstrated the dermal papilla’s key role in hair growth, revealing that it can induce hair formation when implanted into follicles lacking their lower halves- follicles that would otherwise fail to produce hair^32–35^. Later research further emphasized the dermal papilla’s significance in regulating hair shaft properties^36–39^. It also suggested that dermal papilla cells, which are polyclonal and exhibit distinct gene expression profiles compared to dermal fibroblasts, may originate from fibroblasts that have been modified through interactions with follicle epithelial cells^17,40–42^. Moreover, laser ablation of the dermal papilla during the resting phase prevents follicles from re-entering the growth cycle^43^, underscoring its critical role in initiating regeneration.

The current model proposes that the dermal papilla induces hair germ proliferation, while the epithelium forms an intrinsic niche that coordinates the entire hair cycle^12,20,44–50^. However, two major challenges arise with this model. First, inhibiting Wnt protein secretion from dermal papilla cells by targeting the Wnt ligand secretion mediator (*Wls)* gene resulted in only a slight delay in follicle growth, rather than a complete arrest^51^. Second, the transient nature of the hair follicle suggests the need for a ‘long-lived’ cellular source outside the epithelium that interacts with and monitors its state. This hypothetical cell type would integrate local signals and coordinate responses, maintaining the dynamic equilibrium of the epithelial environment.

In light of these questions, we investigated the presence of telocytes in the skin, and their potential role in regulating hair follicle SCs. Our findings demonstrate that telocyte networks form a dynamic, integrative component of the SC niche, playing a central role in coordinating SC activity throughout follicle regeneration.

## Results

### Structural characterization of Foxl1^+^ cells reveals a telocyte network that extends throughout all epithelial layers of the hair follicle

Telocytes have been previously identified through electron microscopy primarily in the human hair follicle, located along the basal bulge and sub-bulge regions, the sites of hair follicle SCs ^52–57^. While telocytes have been suggested to contribute to skin regeneration, this potential function has not been experimentally explored. Given that telocytes in the intestine express the transcription factor Foxl1^7^, we aimed to explore the relationship between this marker and telocyte identity and distribution in the skin.

Foxl1 expression has been documented during hair follicle development and hair placode morphogenesis, particularly within a cluster of mesenchymal cells known as the dermal condensate ^58^. This condensate, which aggregates beneath the developing placode, acts as a signaling hub that regulates follicle morphogenesis and eventually evolves into the dermal papilla^37,58–60^.

To investigate the distribution and structure of Foxl1^+^ cells in the skin, we employed Foxl1-promoter-driven Cre mice crossed with Rosa-mTmG reporter mice^6^, which encode a membrane-bound version of green fluorescence protein (GFP) (**Figure 1A**). This approach provided a detailed visualization of Foxl1^+^ cells.

**Figure 1.**
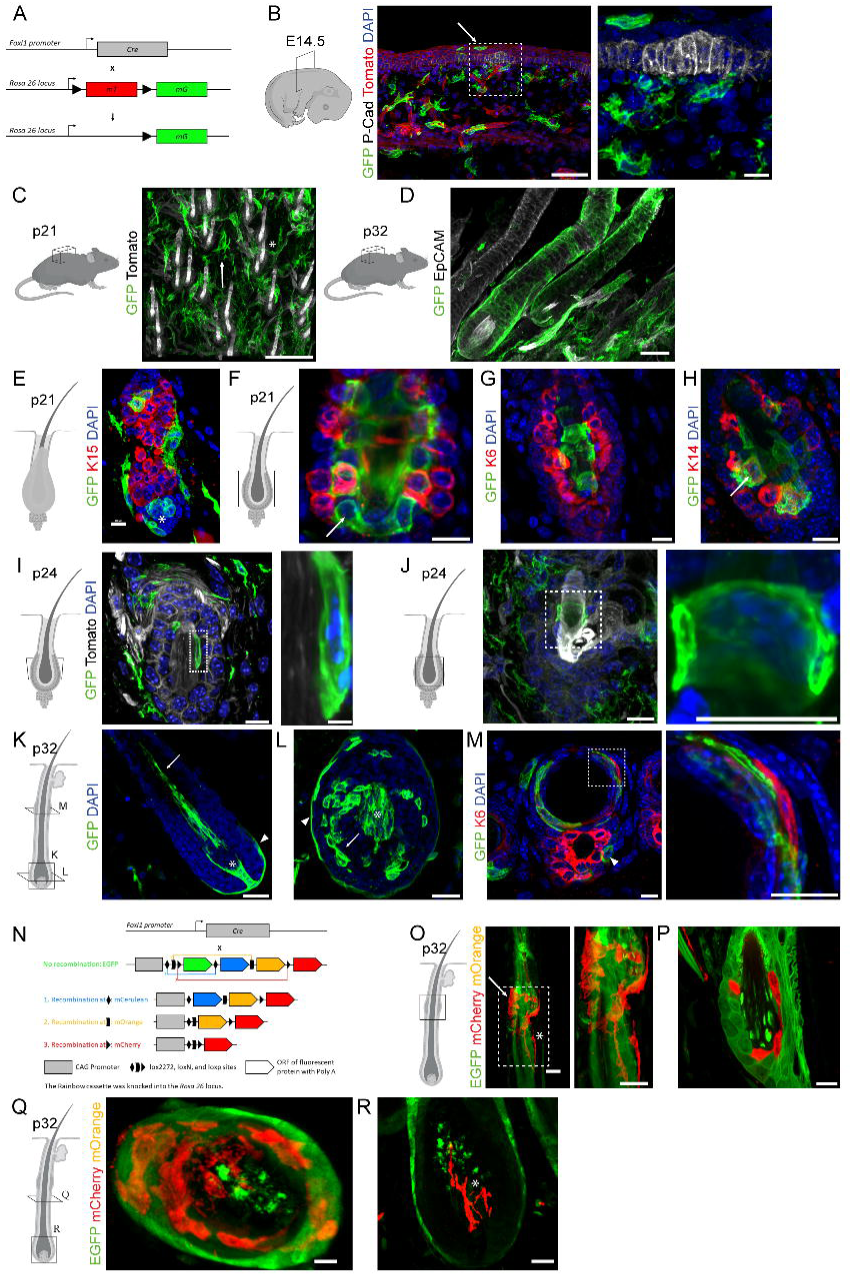
Structural characterization of Foxl1^+^ cells reveals a network that extends throughout all epithelial layers of the hair follicle. (**A**) Scheme of mouse model illustrating how the expression of membrane-bound GFP is driven in Foxl1-Cre-positive cells in the Foxl1-Cre; Rosa-mTmG mouse model. (**B**) Immunofluorescence of E14.5 mouse dorsal skin cross-section reveals the expression of GFP in cells neighboring the developing hair placode stained for P-Cadherin (white). Note the distribution of Foxl1^+^ cells along vessels in close proximity to the placode and scattered along the epidermis (arrow). Boxed region highlights the hair placode region. Scale bar 50μm, 10μm. (**C**) whole-mount immunofluorescence of mouse back skin, following tissue clearing at the hair follicle resting phase on postnatal day 21 (p21), reveals GFP expression in broad cells with multiple extensions (arrow) and thin cells encircling blood vessels (asterisk). Foxl1^+^ cells form a network in close contact with hair follicles visualized in tomato (grey). Scale bar 100μm. (**D**) whole-mount immunofluorescence of p32 mouse back skin following tissue clearing reveals the expression of GFP in cells with a sheath-like morphology that encapsulates the entire hair follicle stained for EpCAM (grey). Scale bar 100μm. (**E-H**) Immunofluorescence of p21 skin sections reveals the expression of GFP in cells distributed within the dermis in close contact with the basal outer bulge and hair germ, beneath the follicle (E asterisk) and along the inner bulge region where no mesenchymal component has been previously reported (F-H). Note the intimate contact Foxl1^+^ cells form with SCs stained for K15 (E-F) or K6 inner bulge epithelial cells (G), as well as with the basal epithelium expressing K14 (H). Arrows indicate nuclei surrounded by GFP signal, but they belong to epithelial cells. Scale bar 10μm. (**I-J**) Cross and longitudinal sections of the bulge region at the onset of growth phase (p24) reveal a single DAPI^+^ cell body associated with GFP signal in the inner bulge (I) and hair shaft (J) regions. Scale bar 10μm, 1μm (I), 10μm (J). (**K-M**) Immunofluorescence at hair follicle full growth phase (p32) reveals GFP expression in cells distributed throughout the follicle: around the follicle surface (K, L, M arrowhead), along the inner concentric layers (L arrow), interspersed between K6-positive differentiated keratinocytes in the companion layer (M boxed region), and along the hair shaft (K arrow). Additionally, GFP expression is noted in the core of the follicle (K, L asterisk). Scale bar 50μm (K-L), 25μm (M). (**N**) Scheme of mouse model illustrating how the expression of membrane-bound Cherry, Orange and Cerulean is driven in Foxl1-Cre-positive cells in the Foxl1-Cre; Rosa-Rainbow mouse model. (**O-R**) Immunofluorescence of longitudinal and cross-sections on p32 mouse back skin reveals the expression of mOrange and mCherry in elongated cells with small cell body and long extensions surrounding the basal outer bulge (O), the inner bulge (O-P), along the epithelial concentric layers (Q) and in the core of the follicle (R asterisk). Note the orientation of Foxl1^+^ cells, positioned horizontally (O arrow) and aligned parallel (O asterisk) to the hair shaft. Scale bar 10μm.

At embryonic day 14.5, during placode development, Foxl1^+^cells appeared with a relatively condensed structure and multiple extensions (**Figure 1B**). These cells were broadly distributed throughout the skin dermis, encircling blood vessels near the developing placode labeled with P-Cadherin (P-Cad). Additionally, Foxl1^+^ cells were detected along the epidermis (**Figure 1B** arrow), a region where, to our knowledge, mesenchymal components have not been previously reported. However, we did not observe clusters of Foxl1^+^ cells beneath the placode, which would typically indicate the dermal condensate.

Considering the physical characteristics of telocytes, particularly their thin membranous cellular extensions, we conducted whole-animal trans-cardiac perfusion (see methods for details) to better preserve cellular structures. To fully visualize the potential 3D network of Foxl1^+^ cells, we performed confocal imaging of immunofluorescent-stained whole mouse back skin after tissue clearing at p21 and p32. At these stages, hair growth is synchronized with the resting (telogen) and full growth (anagen) phases, respectively. 3D image reconstruction revealed an extensive, interconnected network of Foxl1^+^ cells closely associated with hair follicles (**Figures 1C-D)**. During the resting phase, the Foxl1^+^ inter-follicular array consisted of broad cells with multiple extensions (**Figure 1C** arrow) and flat, thin cells encasing blood vessels (**Figure 1C** asterisk). During the growth phase, Foxl1^+^ cells appeared thin and flat and consistently aligned along the basal epithelium in all follicles stained for EpCAM (**Figure 1D**), demonstrating distinct morphological structures that align with the changing architecture of the follicle. It is important to note that the visibility of the GFP signal in these cells depended on the imaging focal plane and fixation protocols. In skin samples from p32 mice that were fixed without perfusion, the GFP signal was undetectable (**Figure S1**), emphasizing the necessity of proper fixation techniques to visualize the delicate structure of Foxl1^+^ cells.

Our findings revealed a network of Foxl1^+^ cells closely associated with hair follicles during regeneration, prompting further investigation into their role in SC activity and differentiation. To explore this, we analyzed the distribution of Foxl1^+^ cells along the concentric epithelial layers of the follicle using longitudinal and cross-sections. Foxl1^+^cells in the dermis were closely in contact with the basal layer of the follicle bulge - home to K15 SCs, as well as the hair germ and were associated with the dermal papilla region (**Figure 1E**, asterisk) - we also detected GFP signals in the inner bulge region (**Figure 1F)**, an area previously believed to lack connective tissue or mesenchymal components. This suggests the presence of mesenchymal elements within the inner layers of the follicle, implying that mesenchymal guidance may extend beyond the dermis.

A detailed analysis, including co-staining with K6 (**Figure 1G)**, a marker of the inner bulge epithelial layer, and K14 (**Figure 1H),** a basal epithelial marker, revealed that Foxl1^+^ cells form a thin sheath covering both the inner and outer bulge (**Figures 1E-H, Videos S1-S2**). The flat, thin structure of Foxl1^+^ cells, along with their extensive interconnected processes and the compact structure of the follicle, can create the impression of cell bodies related to GFP membranes (**Figures 1F and 1H** arrow). However, these nuclei belong to epithelial cells. By correlating the GFP signal with DAPI staining in z-stacks, we found that Foxl1 nuclei are elongated with puncta of heterochromatin, which are typical features of telocytes^61–63^, distinct from the cuboidal epithelial nuclei (**Figures 1I-J, Video S3)**. Within a 20μm thick section, only one GFP related nucleus was observed in the inner follicle regions, indicating that the extensive Foxl1^+^ GFP^+^ network originates from a small number of cells.

During differentiation, we found that the Foxl1^+^ cell network extensively expands across all epithelial layers of the follicle, including inner layers where mesenchymal components were previously thought to be absent. Foxl1^+^ cells were found along the basal outer layer of the follicle (**Figures 1K-M** arrowhead), within the inner bulb and bulge regions (**Figure 1L** arrow**, Video S4)**, along the hair shaft (**Figure 1K** arrow), and intercalated among K6-positive, differentiated keratinocytes in the companion layer (**Figure 1M** boxed region). Foxl1^+^ cells were integrated with or continuous with known mesenchymal components, including those lining the follicle (**Figures 1K-L** arrowhead) and at its core in the bulb region (**Figures 1K-L** asterisk).

Single-molecule RNA fluorescence in-situ hybridization (smFISH) confirmed that all GFP-positive cells derived from Foxl1-Cre, including those within the inner epithelial layers, express Foxl1 mRNA (**Figures S2B-D**), validating that GFP faithfully marks Foxl1^+^ cells. An additional independent inducible Foxl1CreERT2 reporter mouse model^64^ (**Figure S2E**), corroborated these findings, demonstrating similar GFP distribution in the bulge region after tamoxifen induction (**Figures S2F-H**). Immunofluorescence staining for CD31 and Neun confirmed that Foxl1^+^ cells are not endothelial or neuronal in nature but are associated with these cells (**Figures S2I-J**).

To further understand the architecture and orientation of individual Foxl1^+^ cells within the follicle, we used the Rosa-membrane Rainbow reporter mouse line ^65–67^ crossed with Foxl1Cre mice for multispectral marking (**Figure 1N)**. Cells labeled with membrane-bound Orange (orange) or membrane-bound Cherry (red) displayed relatively small cell bodies (**Figures 1O and 1Q)** and elongated extensions, key characteristics of telocytes, aligned parallel to the hair shaft (**Figures 1O and 1R** asterisk) or radially with the concentric epithelial layers (**Figures 1P-Q)**. Fiber-like structures were observed extending horizontally surrounding the basal bulge region (**Figure 1O** arrow) and longitudinally at the core of the follicle bulb (**Figure 1R)**, forming a lattice-like structure.

Thus, Foxl1 marks cells with oval heterochromatin nuclei and elongated processes, forming a robust, interconnected subepithelial network, typical of telocytes. This network integrates with both the basal outer and inner epithelial layers of the follicle, where SCs and their progeny reside. Our findings suggest that the distribution of these mesenchymal cells is more extensive than previously recognized, indicating that hair-inductive capabilities may extend beyond the dermal layer.

### Ablation of Foxl1^+^ telocytes disrupts Wnt-signaling and halts hair follicle growth

To assess the role of telocytes in regulating hair follicle SC activity, we devised a strategy to selectively ablate telocytes at the onset of follicle regeneration. We utilized Foxl1-Cre; Rosa-iDTR mice (Foxl1-iDTR), which express the diphtheria toxin receptor (DTR) in Foxl1^+^ cells through Cre-mediated excision of a stop cassette. This allowed us to specifically target telocytes for ablation following toxin administration.

Given the distinct regenerative demands of hair follicles and intestines - where intestinal SCs are highly proliferative and capable of regenerating the crypt-villus axis from a single cell^68^, while bulge SCs are quiescent and slowly cycling^16,69^ - we hypothesized that partially ablating telocytes would minimally affect intestinal homeostasis, allowing us to focus on their role in hair follicle regeneration.

Ablation was induced at the onset of the hair follicle growth phase (p21-p24), and mice were analyzed the following day to assess immediate effects (**Figure 2A**). smFISH analysis revealed a 70% reduction in Foxl1 mRNA in the bulge of Foxl1- iDTR follicles compared to controls, confirming efficient ablation (**Figure 2B**).

**Figure 2.**
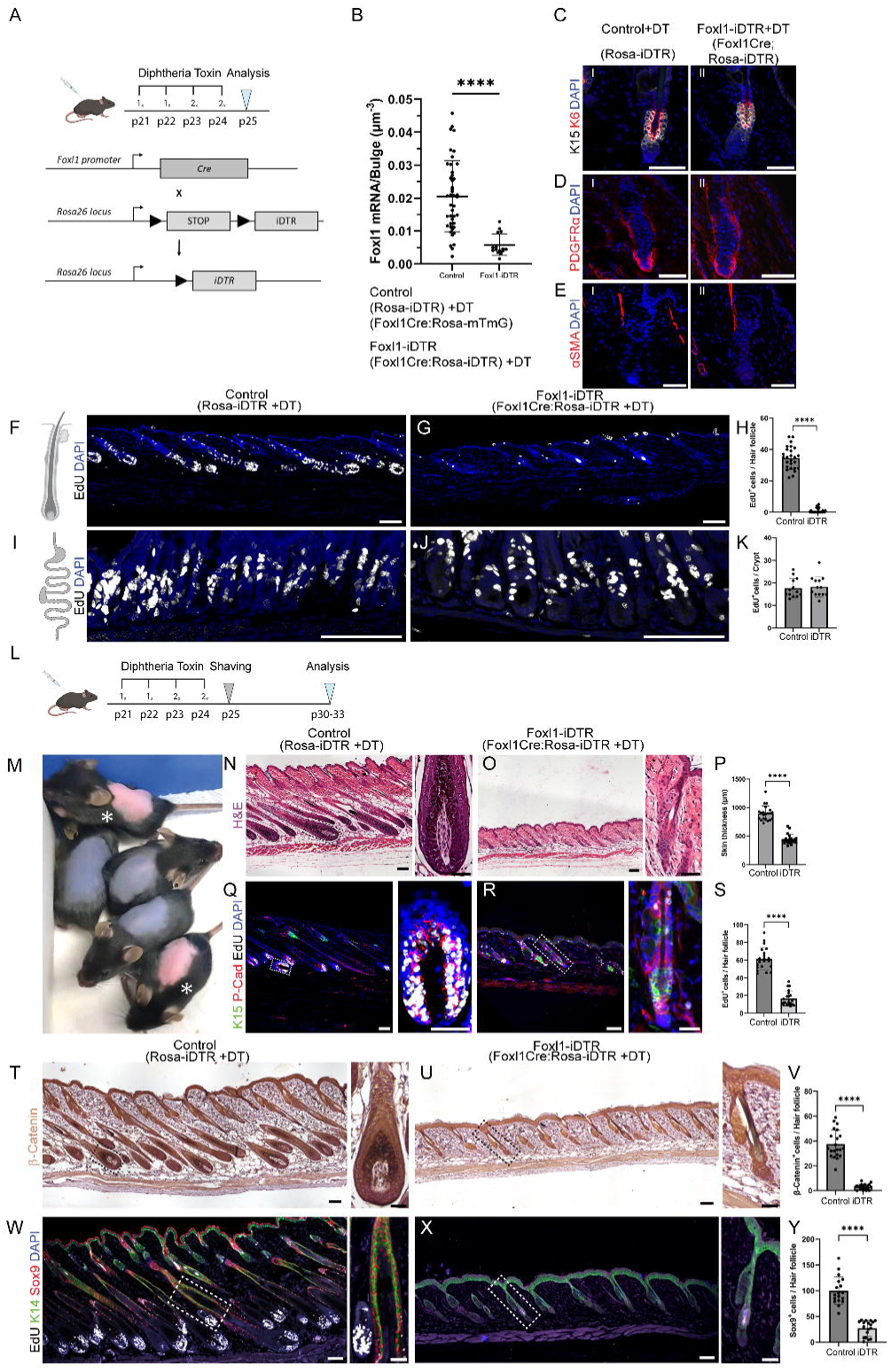
Ablation of Foxl1^+^ telocytes disrupts Wnt-signaling and halts hair follicle growth. (**A-K**) Scheme illustrating the experimental setup where diphtheria toxin induction facilitates partial ablation of telocytes at the onset of growth phase (p21-p24) in Foxl1-Cre-positive cells in the Foxl1-Cre; Rosa-iDTR mouse model. Mice were analyzed at p25 (A). Quantification of Foxl1 mRNA in the bulge region of control mice and Foxl1-iDTR mice (n=3 mice per group <20 follicles for each bar ****P<0.0001, unpaired two-tailed t-test) (B). Note the significant reduction of more than 70% in Foxl1 transcript expression in Foxl1-iDTR mice compared to control mice. Immunofluorescence of control and Foxl1-iDTR follicle longitudinal sections, reveals no differences in the expression and distribution of SCs stained for K15 (white), and K6 (red) inner bulge epithelial cells (CI-II), fibroblasts stained for PDGFRα (DI-II), or arrector pili muscle stained for αSMA (EI-II). Short EdU incorporation (white) in the hair follicle (F-G) and intestinal (I-J) epithelium of control mice (F, I) compared to Foxl1-iDTR mice (G, J). Note the onset of follicle proliferation in control mice, in contrast to the Foxl1-iDTR mice, where proliferation is barely detectable. Intestinal proliferation remains unaffected. Quantification of EdU^+^ cells per follicle (H) or crypt (K) (n=3 mice per group <20 follicles or crypts for each bar, follicles- ****P<0.0001, intestine- no significant difference, unpaired two-tailed t-test). (**L-Y**) Scheme illustrating the experimental setup where diphtheria toxin induced mice were analyzed approximately 10 days after induction, at p30-33 (L). A representative photograph of a litter 10 days following diphtheria-toxin induction (M). Note the difference in back skin appearance: Foxl1-iDTR mice (asterisk) exhibit pink, hairless back skin, while control mice display grey back skin due to pre-emergent hair bristles. H&E staining of skin sections from control mice and Foxl1-iDTR mice (N-P). Note the complete cessation of follicle growth and the dramatic reduction in skin thickness in Foxl1-iDTR compared to control mice. Quantification of skin thickness (P) (n=3 mice per group <20 follicles for each bar ****P<0.0001, unpaired two-tailed t-test). Short EdU incorporation (white) and immunofluorescence for K15 (green) and P-Cadherin (red), in skin longitudinal sections, reveal high EdU incorporation in the bulb region at the bottom, far from the K15-stained stem cells (Q). In contrast, Foxl1- iDTR mice show few EdU-incorporated cells near the bulge region, typical of the onset of growth phase (R). Quantification of EdU^+^ cells per follicle (n=3 mice per group <20 follicles for each bar ****P<0.0001, unpaired two- tailed t-test) (S). Immunohistochemistry for β-Catenin to analyze Wnt activity within the hair follicle. Nuclear β-Catenin staining, indicative of active Wnt-signaling, was dramatically reduced in Foxl1-iDTR mice (U) compared to control mice (T). Quantification of nuclear β-Catenin^+^ cells per follicle (V) (n=3 mice per group <20 follicles for each bar ****P<0.0001, unpaired two-tailed t-test). Immunofluorescence for Sox9 (red), along with K14 (green) and EdU incorporation (white) in control mice (W) and Foxl1-iDTR mice (X). Note the dramatic reduction in Sox9^+^ cells in Foxl1-iDTR follicles compared to control follicles, further supporting impaired Wnt-signaling. Quantification of Sox9^+^ cells per follicle (Y) (n=3 mice per group <20 follicles for each bar ****P<0.0001, unpaired two-tailed t-test). Boxed regions highlight the follicle structure in each group. Scale bar 100μm; boxed regions 25 μm.

We examined key markers of the bulge (K15 and K6), surrounding stroma (PDGFRα) and arrector pili muscle (αSMA). No differences were observed between Foxl1-iDTR mice and controls (**Figures 2C-E**), indicating that Foxl1^+^ cell ablation specifically affected telocytes without disrupting other Foxl1-negative stromal cells or K15 / K6 – expressing cells in the bulge. While intestinal regeneration remained intact (**Figures 2I-K**), remarkably, hair follicle regeneration was completely halted in Foxl1-iDTR mice, as shown by EdU labeling of S-phase cells (**Figures 5F-H**). This indicates that, upon telocyte ablation, hair follicle proliferation and progression into the growth phase were entirely blocked.

We analyzed this striking phenomenon in further detail. At the peak of growth phase (p30-33) (**Figures 2L-Y**), control mice displayed pre-emergent hair bristles, whereas Foxl1-iDTR mice exhibited hairless pink skin (**Figure 2M**). Histological analysis revealed a complete halt in hair follicle growth in telocyte-deficient mice, with follicles remaining in the resting phase, while control mice showed elongated follicles with new hair shafts (**Figures 2N-O**). Skin thickness in the Foxl1-iDTR mice was halved compared to controls (**Figure 2P**), and EdU labeling with K15 staining showed significantly reduced stem and progenitor cell proliferation in Foxl1-iDTR mice (**Figures 2Q-S**). In controls, K15-expressing SCs were localized in the upper region of the follicle (**Figure 2Q**), characteristic of the growth phase, while in Foxl1-iDTR mice, K15 SCs were confined to the follicle base, with minimal proliferation in the hair germ (**Figure 2R**).

Since hair follicle SC proliferation is Wnt-dependent ^26–28,70–72^, we hypothesized that the absence of follicle growth in Foxl1-iDTR mice was due to disrupted Wnt signaling. Indeed, we observed a significant reduction in nuclear β-catenin, an indicator of active Wnt signaling, in Foxl1-iDTR follicles, confirming a loss of canonical Wnt signaling (**Figures 2T-V**).

We also examined Sox9, a pioneer factor^73^ that relies on Wnt gradients for SC specification^48^. Following diphtheria toxin treatment, K15 expression persisted in the follicles of Foxl1-iDTR mice, but the number of Sox9-expressing cells was dramatically reduced compared to controls (**Figures 2W-Y**). This dramatic decrease underscores the impaired Wnt signaling activity and suggests a strong association between Sox9 and SC activity. These results indicate the crucial role of telocytes in inducing hair follicle regeneration.

### Molecular characterization of Foxl1^+^ telocytes reveals their role in follicle-instructive mesenchyme

To better understand the relationship of telocytes with known follicle-instructive mesenchymal components, we further analyzed publicly available RNA sequencing data from fetal mouse skin samples. The analysis confirmed the presence of Foxl1 mRNA in mesenchymal components^58,74–77^, supporting the mesenchymal identity of these cells. Specifically, we examined data from E15.0 fetal mouse skin ^76^, which detailed the classification of instructive mesenchymal components involved in hair placode morphogenesis. During this stage, mesenchymal cells cluster beneath the developing placode to form the dermal condensate, a precursor to the dermal papilla that expresses Sox2. The mesenchymal populations were classified as dermal condensate cells in the formation stage (DC1), those in the polarized downgrowth hair germ stage (DC2), unclustered progenitors (Pre-DC), Schwann cells lining nerve projections, fibroblasts, and PDGFRα^-^/Sox2^-^ negative cells.

Bulk RNAseq revealed that Foxl1 expression was highest in both dermal condensate populations (DC1 and DC2) and in progenitor pre-DC cells (**Figure 3A**). Foxl1 was also present in Schwann cells and PDGFRα^-^/ Sox2^-^ negative cells, while its expression was minimal to absent in PDGFRα^High^ fibroblasts. This suggests that Foxl1^+^ cells have a distinct molecular signature, differentiating them from fibroblasts and aligning them more closely with the characteristics of telocytes- signaling hub cells with unique traits and a clearly defined identity separate from fibroblasts^1,4,61,63,78^.

**Figure 3.**
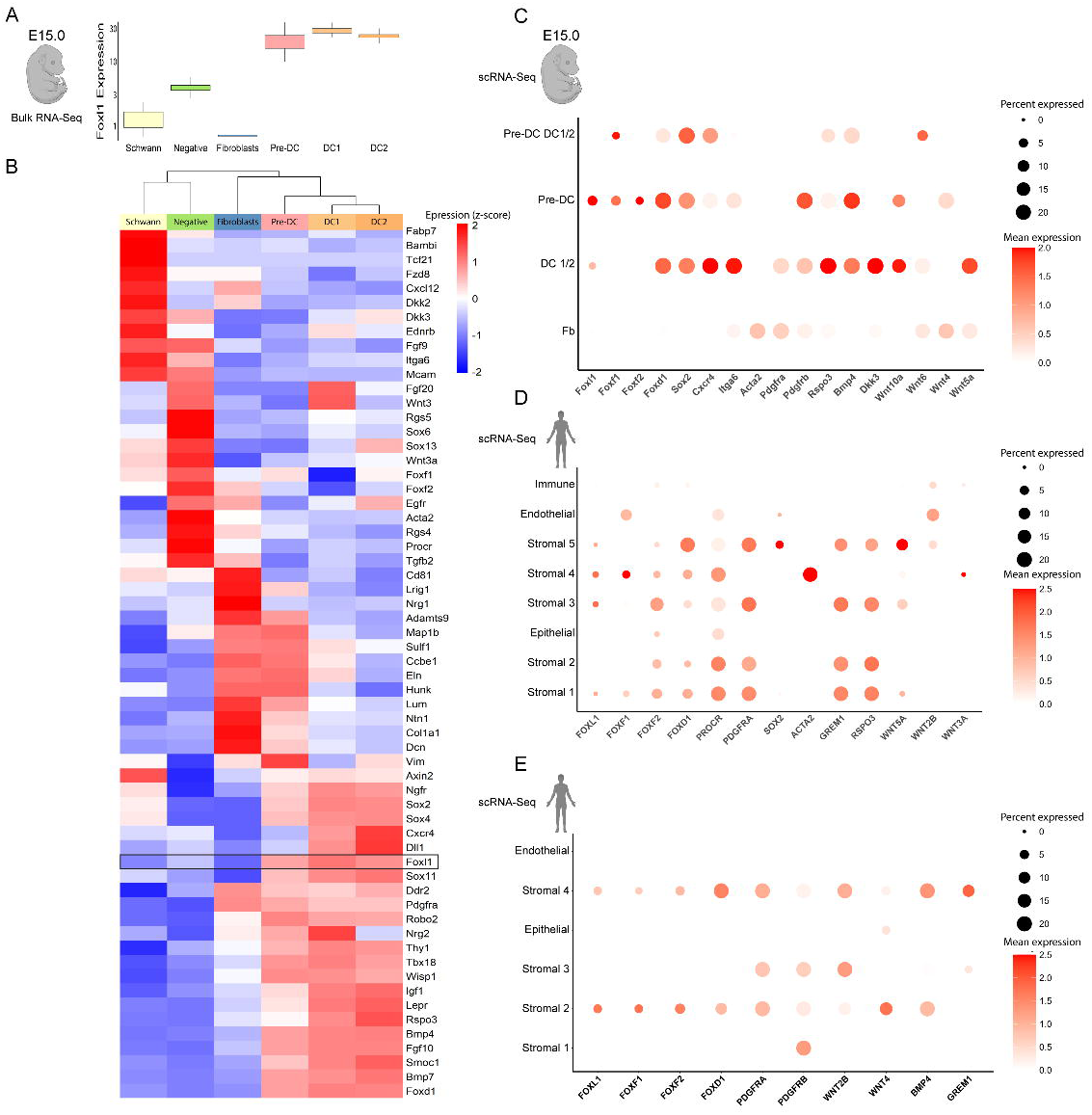
Molecular characterization of Foxl1^+^ telocytes reveals their role in follicle-instructive mesenchyme. (**A-B**) Gene expression from bulk RNA-Seq data of E15.0 mouse skin, from six cell subtypes FACS sorted based on Sox2-EGFP expression. DC1- Sox2^High^/CXCR4^Low^ dermal condensate cells during their formation stage, DC2- Sox2^High^/CXCR4^High^ dermal condensate cells at the polarized downgrowth hair germ stage, Pre-DC- Sox2^Low^/PDGFRα^Low^/Itgα6^Low^ unclustered progenitors of dermal condensate cells, Schwann cells - Sox2^Low^/PDGFRα^Low^/Itgα6^High^, Fibroblasts- PDGFRα^High^, Negative- PDGFRα^-^/Sox2^-^. Boxplot showing the interquartile range of Foxl1 normalized expression (DESeq2, normalized counts), across the six subtypes (A). Note the minimal to absent expression of Foxl1 in fibroblasts. Row normalized (z score) heatmap showing selected differentially expressed genes from each subtype (B). (**C-E**) DotPlots showing the mean normalized expression (color, log [TP10K+1]) and percent of cells expressing (dot size) from scRNA-Seq expression data. Single cell profiles from each study were analyzed and re-clustered using an updated standardized analysis pipeline (R, Seurat), cell subtype clusters were identified by Leiden graph clustering, and annotated using marker genes and prior sub-type annotations. E15.0 mouse skin showing two Foxl1^+^ clusters identified as Pre-DC and DC1/2 (C). Data from human skin showing FOXL1 expressing cells across diverse stromal clusters enriched for key signaling molecules (D-E).

Transcriptome analysis of Foxl1^+^ cells revealed a molecular signature that links them to mesenchymal cells with niche-supporting functions, interacting with both developing placode epithelia and neurons (**Figure 3B**). Notably, this analysis identified a previously unrecognized population with high expression of Foxf1 and Foxf2, transcription factors involved in Sonic Hedgehog signaling and intestinal development ^79–81^. This population also expressed Procr and Mcam, markers linked to intestinal telocytes (Gharbi *et al*., work in progress). These findings suggest that Foxl1^+^ cells in both the skin and intestine may share common markers and potentially analogous functions. Moreover, this may indicate that these markers correspond to the previously unrecognized population of telocytes within the inner layers of the hair follicle. Foxl1^+^ cells were particularly enriched in signaling molecules essential for both follicle and intestinal regeneration, including Wnt, Rspondin, Bmp, Fgf, Igf, Notch, Tgfβ, Lepr and Egfr ^22,82–97^, reinforcing the role of telocytes as central signaling hub cells.

To further characterize the diversity of Foxl1^+^ cells, we analyzed expression profiles of fetal mouse skin captured by single-cell RNAseq, focusing on fibroblasts, pre-DC, DC1 and DC2 clusters, which excluded Schwann cells and PDGFRα^-^/ Sox2^-^ negative cells ^76^. Out of 1,604 cells analyzed, only 23 showed Foxl1 expression, highlighting the difficulty in detecting lowly expressed transcripts through single-cell RNAseq. This also suggests that optimizing tissue dissociation protocols may be crucial for obtaining viable Foxl1^+^ cells. Nevertheless, Foxl1^+^ cells were identified within the pre-DC, DC1 and DC2 clusters but were absent in fibroblasts (**Figure 3C**), aligning with bulk RNAseq findings. Interestingly, the Pre-DC Foxl1 cluster also expressed both Foxf1 and Foxf2, indicating that in the skin, these factors are co-expressed in Foxl1^+^ cells beyond the PDGFRα^-^/Sox2^-^ negative population and may serve as potential markers for telocytes.

To strengthen our observations from developing mouse skin, we analyzed data from human skin in two independent studies ^98,99^ to identify common markers shared across different species that can be used to identify telocytes. FOXL1 expression was identified exclusively in stromal cell populations from both studies, highlighting the mesenchymal origin of these cells (**Figures 3D-E**). Further investigation revealed that FOXL1 subpopulations expressed mesenchymal markers such as PDGFRA, PDGFRB and ACTA2. Additionally, some FOXL1 subpopulations exhibited expression of FOXF1, FOXF2 and the dermal condensate markers SOX2 and FOXD1, aligning well with our observations in the intestine and fetal mouse skin. Furthermore, FOXL1^+^ cells demonstrated expression of signaling molecules potentially involved in supporting SC activity, including Wnt ligands (Wnt2b, Wnt3a, Wnt4 and Wnt5a), the Wnt agonist Rspondin 3 and the Bmp-inhibitor Grem1 (**Figures 3D-E**). These factors likely explain the critical role of telocytes in promoting hair follicle regeneration.

Altogether, this analysis suggests that across species, Foxl1^+^ cells are enriched in signaling molecules, expressing common transcription factors such as Foxf1, Foxf2, Sox2 and Foxd1, which may be linked to telocyte identity.

### Foxl1^+^ telocytes retain consistent markers through both placode morphogenesis and follicle regeneration

Next, we explored the association between telocytes and the markers identified in Foxl1^+^ cells from the RNAseq analysis by performing immunostaining on skin sections from Foxl1 reporter mice. During placode morphogenesis, at E14.5, while Sox2 and Ngfr clearly labeled aggregates of cells at the base of the developing placode, characteristic of the DC (**Figures 4B-C**), these cells were GFP-negative. Some telocytes did express Sox2 but were positioned adjacent to, rather than within, the DC (**Figure 4B)**. Notably, during the downgrowth hair germ stage, telocytes were observed within the inner layers of the nascent follicle (**Figure 4C** arrow), suggesting their early incorporation into these layers during follicle morphogenesis.

**Figure 4.**
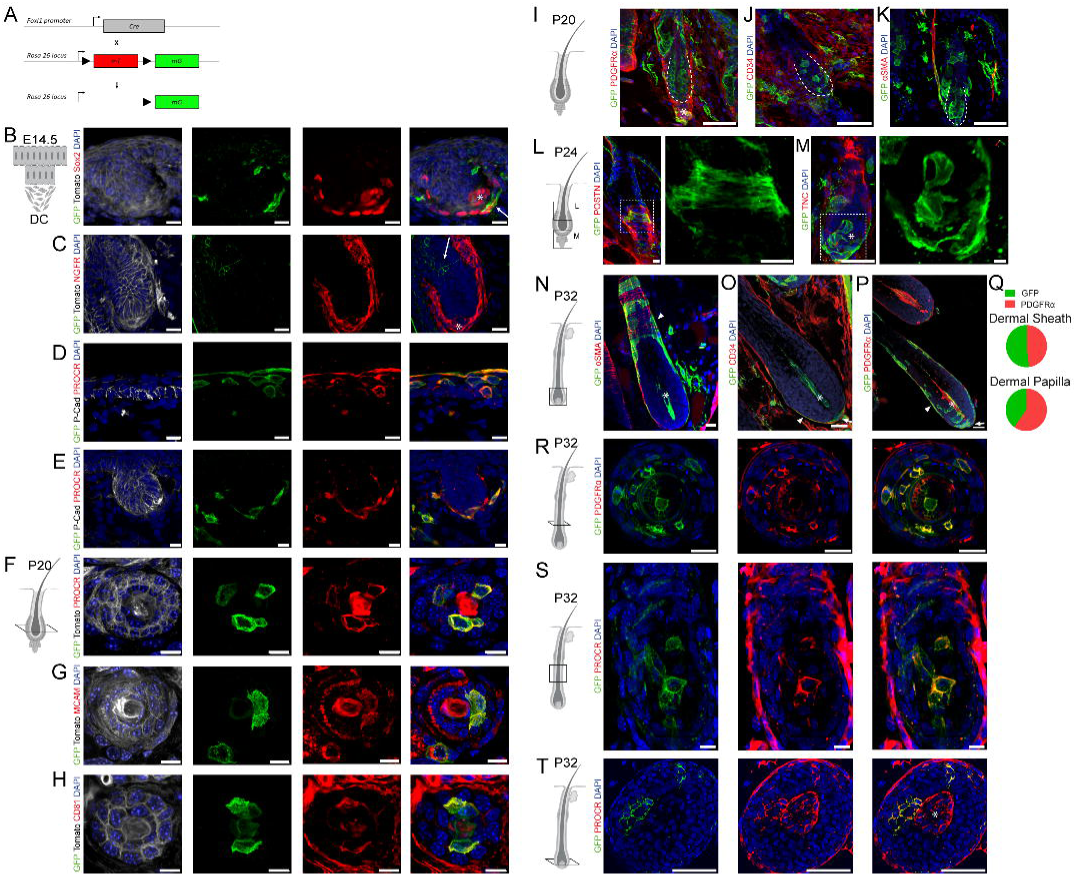
Foxl1^+^ telocytes retain consistent markers through both placode morphogenesis and follicle regeneration. (**A**) Scheme of mouse model illustrating how the expression of membrane-bound GFP is driven in Foxl1-Cre-positive cells in the Foxl1-Cre; Rosa-mTmG mouse model. (**B-C**) Immunofluorescence of E14.5 mouse dorsal skin cross-sections reveals the expression of GFP in cells expressing Sox2 (B arrow) surrounding but not within the dermal condensate, which is stained for Sox2 and Ngfr (B-C asterisk). Note the expression of GFP in cells along the inner layers of the developing hair placode (C arrow). Scale bar 10μm. (**D-E**) Immunofluorescence of E14.5 mouse dorsal skin cross-sections reveals that Foxl1^+^ cells (green) along the epidermis and surrounding the placode are positive for Procr (red). Scale bar 10μm. (**F-H**) Immunofluorescence of p20 hair follicle cross-sections reveals that Foxl1^+^ cells along the inner bulge region are positive for Procr (F red), Mcam (G red) and Cd81 (H red). Scale bar 10μm. (**I-K**) Immunofluorescence of p20 hair follicle longitudinal sections reveals that Foxl1^+^ cells (green) within the dermis, but not along the bulge region (dashed line), are stained positive for PDGFRα and Cd34. Note the expression of PDGFRα in Foxl1^+^ cells beneath the follicle indicating that these cells belong to the dermal papilla component (I asterisk) and the close association of Foxl1^+^ cells with the arrector pili muscle stained for αSMA (K red). Scale bar 50μm. (**L-M**) Immunofluorescence of p24 hair follicles at the transition from the resting phase to the growth phase reveals distinct structure of Foxl1^+^ cells embedded within the basement membrane stained for Postn (L red) and Tnc (M red). Around the bulge region, Foxl1^+^ cells displayed fiber-like structures stretching horizontally around the follicle (L). At the base, Foxl1^+^ cells exhibited a thin, flat morphology along the basal outer follicle layer in continuous with the dermal papilla region (M). Scale bar 10μm. (**N-P**) Immunofluorescence of p32 skin sections at the follicle full growth phase reveals that Foxl1^+^ cells (green) surrounding the follicle, but not in the core region (N asterisk), are positive for αSMA (N red), indicating that these cells are part of the dermal sheath component. Some cells at the base of the follicle, but not in the core region (O asterisk) are positive for Cd34 (O arrow) and PDGFRα (P arrow), indicating these cells are part of the dermal cup cell component. Foxl1^+^ cells at the core of the follicle are stained positive for PDGFRα (P asterisk), indicating these cells are part of the dermal papilla component. Scale bar 50μm. (**Q**) Quantification of the GFP^+^/PDGFRα^+^ signal ratio per follicle, presented as a percentage, shows a mean of 52% ± 27 in the dermal sheath and 40% ± 22 in the dermal papilla (n=3 mice, 150 follicles for each chart). ® Immunofluorescence of p32 hair follicle cross- section reveals that Foxl1^+^ cells (green) along the hair shaft are positive for PDGFRα (R red). Scale bar 25μm. (**S**) Immunofluorescence of p32 hair follicle longitudinal section reveals that Foxl1^+^ cells in the inner follicle layers above the bulb region are positive for Procr. Scale bar 10μm. (**T**) Immunofluorescence of p32 hair follicle cross- sections reveal that Foxl1^+^ cells in the inner layers of the bulb region are stained positive for Procr. Asterisk-dermal papilla. Scale bar 50μm.

To further investigate telocytes that do not express Sox2, we stained for Procr, which marked a PDGFRα^-^/Sox2^-^ negative population in the fetal skin. Procr-labeled telocytes were scattered along the epidermis **(Figure 4D)** and in the dermis (**Figure 4E)**, adjacent to the developing placode, indicating potential roles in epidermal differentiation and placode development.

During follicle regeneration at the resting phase (p20), inner bulge telocytes expressed Procr, Mcam and Cd81 (**Figures 4F-H**), which were identified in a PDGFRα^-^/Sox2^-^ negative cluster and Schwann cells in fetal mouse skin. In contrast, telocytes throughout the dermis expressed dermal markers such as PDGFRα and Cd34 (**Figures 4I-J)**, highlighting the diversity within telocyte populations. Telocytes located beneath the follicle were positive for PDGFRα, indicating their involvement in the dermal papilla component (**Figure 4I)**. At this stage, telocytes were negative for α smooth muscle actin (αSMA) but showed close association with the αSMA^+^ arrector pili muscle (**Figure 4K**).

During the transition from the resting phase to the growth phase, (p24 Anagen II)^100^, peripheral telocytes in the bulge exhibited low αSMA staining (data not shown) and displayed fiber-like structures extending horizontally around the follicle within the extracellular matrix stained for periostin (POSTN) (**Fig. 4L).** Conversely, telocytes at the follicle base had a thin, flat morphology along the basal outer follicle layers, continuous with the dermal papilla region, and were embedded in the extracellular matrix stained for tenascin (TNC) (**Fig. 4M)**. These findings suggest dynamic changes in telocyte structure during the transition from resting to growth phases.

During the growth phase, the expanding follicle is covered by a dermal sheath expressing αSMA and PDGFRα^75^. The dermal sheath has demonstrated inductive roles, capable of restoring follicle growth when implanted under follicles where the lower halves were removed^101^. At the full growth phase (p32), peripheral telocytes surrounding the follicle’s surface expressed αSMA and PDGFRα (**Figures 4N and 4P**), indicating their role in the dermal sheath. A subset of telocytes located in the dermal cup at the follicle’s base expressed Cd34 and PDGFRα (**Figures 4O-P**), with only a fraction of PDGFRα−expressing dermal papilla cells being GFP-positive (**Figure 4P**). Quantification revealed that telocytes represented approximately 50% of the dermal sheath and 40% of the dermal papilla compartments (**Figure 4Q**). Cross-sections above the follicle bulb showed that telocytes in contact with the inner follicle expressed PDGFRα at lower levels compared to the dermis (**Figure 4R**). Procr, which marks telocytes in the inner bulge, also labels telocytes in the inner bulb (**Figures 4S-T**). This suggests that the previously unrecognized PDGFRα^-^/Sox2^-^/Procr^+^ cell population identified during placode morphogenesis corresponds to telocytes distributed within the inner bulge and bulb layers during follicle regeneration.

Across various systems - fetal mouse placode morphogenesis, human skin, and mouse follicle regeneration - telocytes exhibit mesenchymal origins with diverse marker expression. Despite this diversity, telocytes consistently express common markers such as PDGFRα, αSMA, Cd34, Procr, Mcam, Foxf1, Foxf2, Foxd1 and Sox2, and are notably enriched in signaling molecules.

### Foxl1^+^ telocytes compartmentalize mRNA molecules for localized, phase-specific signaling

The significance abundance of signaling molecules in telocytes led us to investigate whether their expression changes dynamically according to follicle compartment and the stage of regeneration. We focused on the Wnt and Bmp pathways, which have opposing roles in SC activity. Canonical β-catenin-dependent Wnt activity first emerges in progenitor hair germ cells during the late resting phase^17^. As growth begins, this activity extends to both hair germ and bulge SCs^31^. By the full growth phase, SCs return to quiescence^12^, with Wnt activity becoming confined to transit-amplifying cells in the follicle bulb^46^, following proliferating regions. In contrast, Bmp signaling maintains SC quiescence^12^ and supports differentiation into hair^102^, peaking in the bulge during the resting phase before tapering off toward its end^103,104^.

We analyzed follicle regeneration at three key stages: late resting phase (p20), early growth (p24 Anagen II), and full growth (p30). Telocytes were classified by their distribution across different follicle compartments: dermal papilla (DP), hair germ (HG), bulge and isthmus - an area above the bulge with a stem cell population contributing to skin lineages, including sebaceous gland, interfollicular epidermis and hair^105^. By the full growth phase, telocytes in the bulb were further divided into outer bulb (OB) telocytes, which contact the basal outer epithelium, and inner bulb (IB) telocytes, which interact with the inner epithelial layers.

To assess telocyte involvement in the niche, we used skin sections of Foxl1 reporter mice to label telocytes with GFP, and we performed smFISH to detect mRNAs encoding niche factors critical for stem cell activity. We quantified the number of mRNA molecules associated with the GFP signal in each compartment (**Figure 5A**).

**Figure 5.**
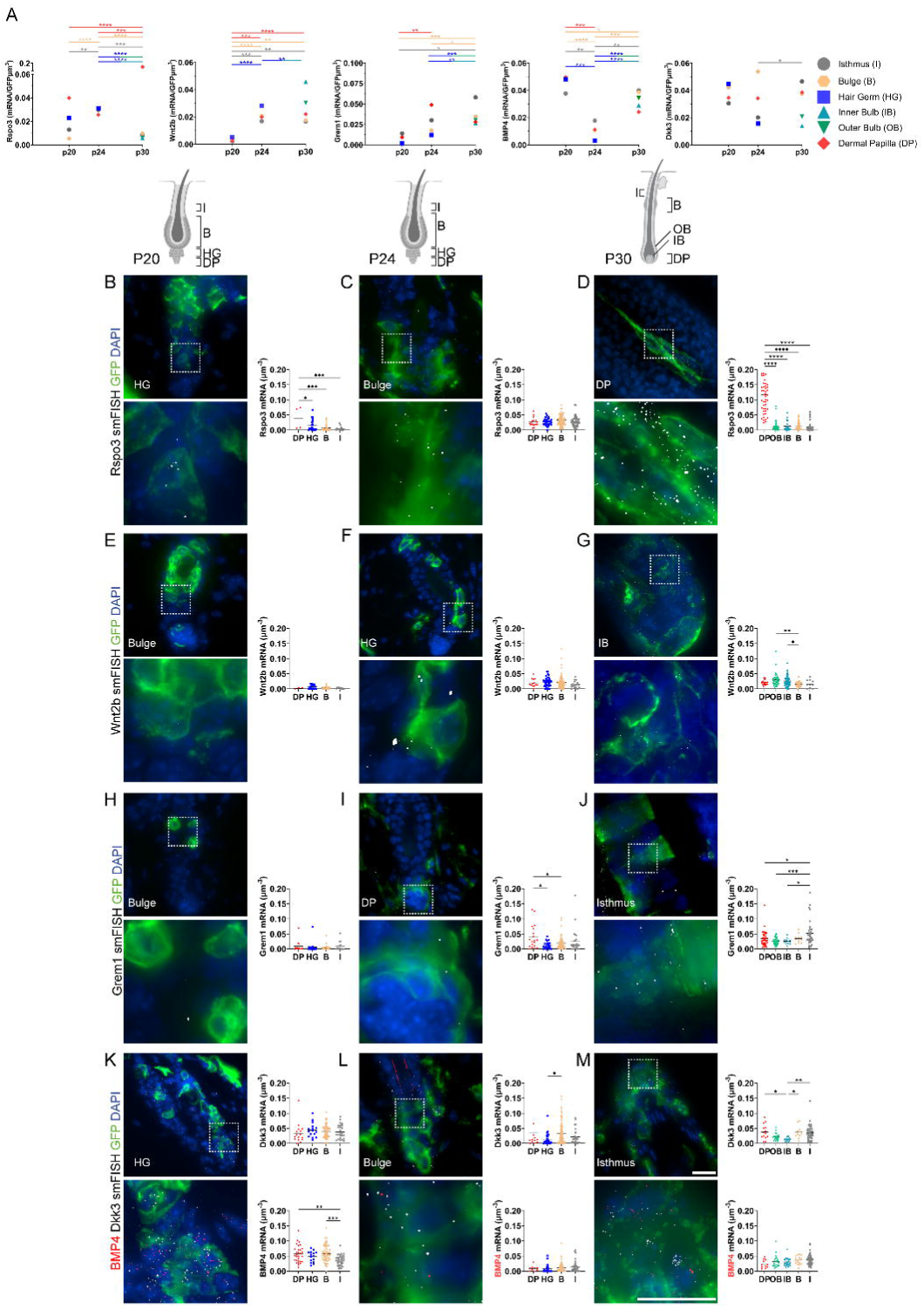
Foxl1^+^ telocytes compartmentalize mRNA molecules for localized, phase-specific signaling. (**A**) Quantification of mRNA molecules encoding signaling proteins per GFP signal during follicle regeneration at p20, p24 and p30, across different follicle regions: the Isthmus (I), Bulge (B), Hair Germ (HG), Inner Bulb (IB), Outer Bulb (OB) and Dermal Papilla (DP). Note the early expression of Rspo3 in telocytes in the DP and HG regions, already detectable by the end of the resting phase (p20). As the growth phase begins (p24), both Rspo3 and Wnt2b show increased expression in all telocyte components, with Rspo3 in the dermal papilla and Wnt2b localized in the inner bulb by the full growth phase (p32). (n=3 mice per group <20 follicles for each bar ****P<0.0001, ordinary one-way ANOVA) (**B-D**) smFISH images display longitudinal and cross-sections of p20, p24 and p30 follicles, stained for GFP to label telocytes and hybridized to detect Rspo3 mRNA (grey dots). At the end of the resting phase, Rspo3 is detected in HG telocytes (B), its expression increases in all telocyte components at the onset of growth phase (p24) (C). By the full growth phase (p30), Rspo3 expression localizes primarily to the dermal papilla, showing a tenfold increase compared to p20 (A, D). (**E-G**) smFISH images display longitudinal sections of p20, p24 and p30 follicles, hybridized to detect Wnt2b mRNA (grey dots). At the onset of growth phase (p24), Wnt2b expression increases (F). By p30, high Wnt2b expression is observed in telocyte extensions within the inner bulb (G), where proliferating progenitors reside. (**H-J**) smFISH images display longitudinal sections of p20, p24 and p30 follicles, hybridized to detect Grem1 mRNA (grey dots). Grem1 is expressed in the dermal papilla at the onset of growth phase (p24) (I) and further increases along the isthmus by the full growth phase (p30) (J). (**K-M**) smFISH images show longitudinal sections of p20, p24 and p30 follicles, hybridized to detect Bmp4 (red) and Dkk3 (white) mRNAs. During the resting phase (p20), high expression of both Bmp4 and Dkk3 is observed in all telocyte compartments (K). At the onset of the growth phase (p24), Bmp4 expression decreases across all regions, while Dkk3 decreases in the HG and Isthmus but remain high in the bulge (L). By the full growth phase (p30), both transcripts show increased expression in the isthmus region compared to their levels at the onset of growth (p24) (M). Scale bar 10μm.

Rspo3, a key inducer of Wnt signaling that regulates the duration of the hair follicle growth phase^106^, is lowly expressed but highly concentrated in telocytes within the HG and DP compartments at the end of the resting phase (**Figure 5B**). This coincides with the early activation of Wnt signaling in the hair germ epithelium. As growth begins, Rspo3 expression increases in telocytes across all follicle compartments (**Figures 5B-C**), with the highest levels observed in telocytes along the bulge and hair germ, where stem and progenitor cells are actively proliferating. By the full growth phase, when Wnt activity and epithelial proliferation are confined to the follicle bulb, Rspo3 expression becomes specifically localized to dermal papilla telocytes, showing more than a tenfold increase compared to the resting phase (**Figure 5D**).

We then examined Wnt2b, which signals through the canonical Wnt/β-catenin pathway. At the end of the resting phase, only a few sparse Wnt2b mRNA molecules were detected in the bulge and HG compartments (**Figure 5E**). As the growth phase progressed, Wnt2b expression increased in telocytes, with the highest – though not significantly different– levels observed in the HG region (**Figure 5F**). By p30, Wnt2b mRNA peaked in telocytes located along both the outer and inner bulb epithelium (**Figure 5G**), aligning with regions of active Wnt signaling in the follicle.

Since canonical Wnt signaling in hair germ progenitors coincides with Bmp inhibition^102,103,107,108^, we also examined the expression of Gremlin1 (Grem1), a Bmp inhibitor. Grem1 mRNA levels steadily increased in telocytes across the bulge, HG, OB and IB from the resting phase through to full growth (**Figure 5A**). By p24, the highest Grem1 expression was observed in dermal papilla telocytes, and by p30, Grem1 peaked in the isthmus region, corresponding with elevated levels of Bmp4 and the Wnt inhibitor Dickkopf3 (Dkk3) (**Figure 5M**), suggesting a potential role in modulating Bmp activity. Notably, the expression patterns in the isthmus during various stages closely mirrored those in the bulge, indicating potential coordination between these regions via telocyte signaling.

Conversely, Bmp4 and Dkk3 mRNA levels were elevated at p20. Bmp4 expression decreased by p24, showing the lowest levels in HG telocytes, while Dkk3 remained high in bulge telocytes, potentially modulating Wnt signaling. By p30, Dkk3 expression had decreased in bulge telocytes and was lowest in the bulb region.

In conclusion, telocytes compartmentalize mRNAs for Wnt and Bmp signaling factors in a spatially and temporally regulated manner, facilitating localized, phase-specific signaling. This suggests a key role for telocytes as a source of signals that orchestrate hair follicle regeneration.

### Hair follicle regeneration depends on Wnts secreted from Foxl1^+^ telocytes

To determine whether Wnt signals expressed by telocytes are functional and play a role in inducing hair follicle SC activity, we generated Foxl1CreERT2; Porcn^Δ/Y^ mice (Foxl1-PorcnΔ) by crossing inducible Foxl1CreERT2 mice^64^ with mice carrying a floxed allele of the X-linked *Porcn* gene^109^. This model allowed us to conditionally ablate Wnt protein secretion specifically from telocytes.

To minimize intestinal injury, we administered a lower dose of tamoxifen than is typically used for labeling telocytes. Induction took place at p21-23 and skin analysis was conducted during full growth phase at p30-33 (**Figure 6A**). Foxl1-PorcnΔ mice showed a slight reduction in weight gain starting three days post-tamoxifen induction compared to controls, but the difference was minor and statistically insignificant (**Figure 6B**).

**Figure 6.**
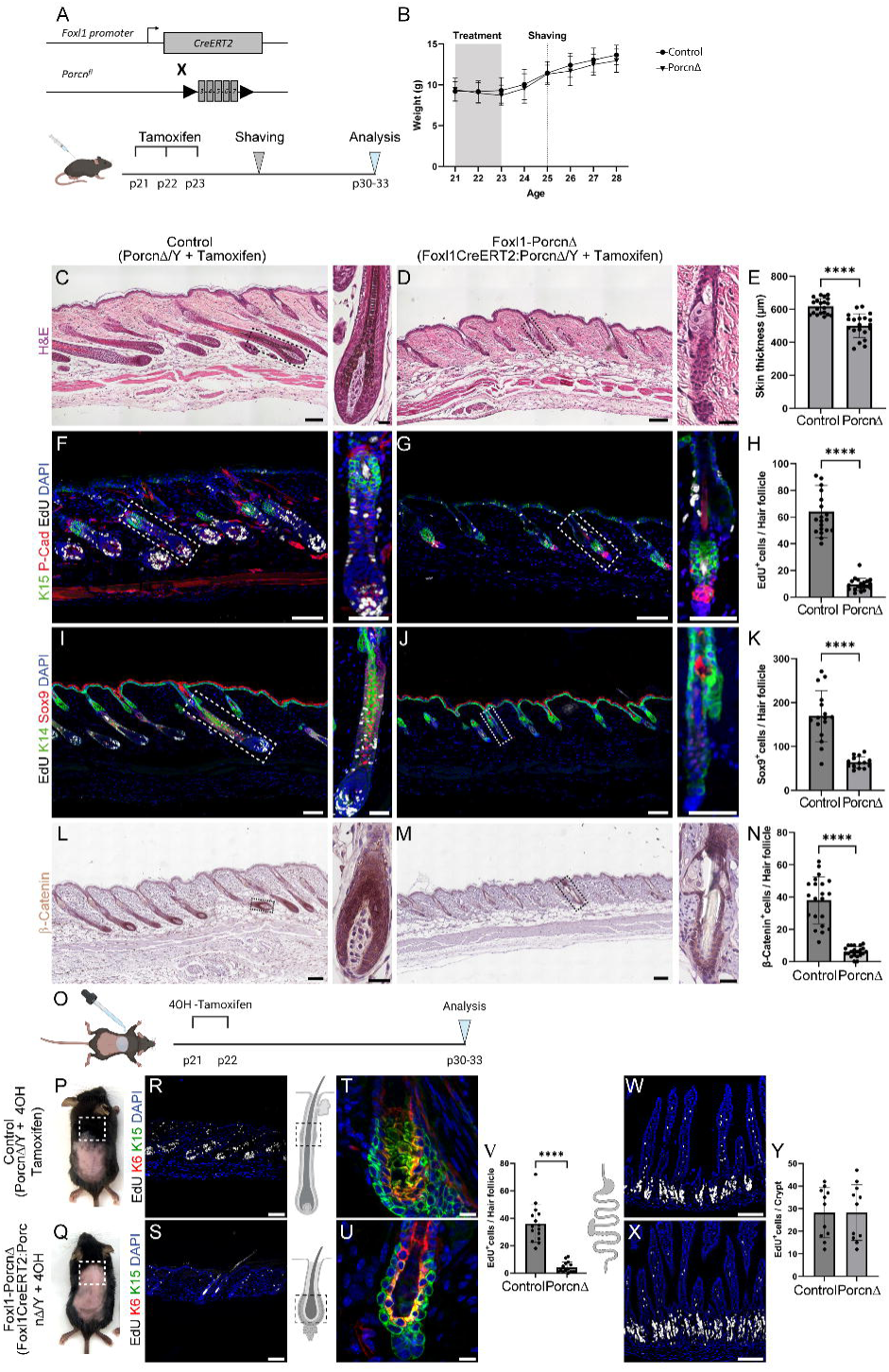
Hair follicle regeneration depends on Wnts secreted from Foxl1^+^ telocytes. (**A**) Scheme illustrating the experimental setup where tamoxifen induce ablation of Wnt secretion from telocytes at the onset of growth phase (p21-p23) in Foxl1-Cre- positive cells in the Foxl1-CreERT2; Porcn^Δ/Y^ mouse model. Skin was analyzed at p30-33. (**B**) Weight curve of Porcn^Δ/y^ (Control n=7) and Foxl1CreERT2; Porcn^Δ/y^ mice, (Foxl1-PorcnΔ n=10) induced with tamoxifen and weighed daily, shows no statistically significant difference in weight between Foxl1-PorcnΔ and control mice (repeated measurements two-way ANOVA). (**C-E**) H&E staining of skin sections from p30-33 mice comparing control mice (C) and Foxl1-PorcnΔ mice (D). Note the cessation of follicle growth in Foxl1-PorcnΔ mice compared to control mice. Quantification of skin thickness (E) (n=3 mice per group <20 follicles for each bar ****P<0.0001, unpaired two-tailed t-test). (**F-H**) Short EdU labeling (white) and immunofluorescence for K15 (green) and P-Cadherin (red) in skin longitudinal sections reveal high EdU incorporation in cells located at the bottom of the follicle, at the bulb region, in control mice compared to the limited EdU incorporation to the hair germ region in Foxl1-PorcnΔ mice. Quantification of EdU^+^ cells per follicle (H) (n=3 mice per group <20 follicles for each bar ****P<0.0001 unpaired two-tailed t-test). (**I-K**) Immunofluorescence for Sox9 (red), K14 (green) along with EdU labeling (white) in control mice (I) compared to Foxl1- PorcnΔ mice (J). Quantification of Sox9^+^ cells per follicle (K) (n=3 mice per group <20 follicles for each bar ****P<0.0001, unpaired two-tailed t-test). Note the reduction in Sox9^+^ cells in Foxl1-PorcnΔ compared to control follicles, supporting impaired Wnt-signaling. (**L-N**) Immunohistochemistry for β-Catenin to analyze Wnt activity within the follicles. Reduced nuclear β-Catenin staining is observed in Foxl1-PorcnΔ (M) compared to control (L) mice further supporting impaired Wnt-signaling. Quantification of nuclear β-Catenin^+^ cells per follicle (N) (n=3 mice per group <20 follicles for each bar ****P<0.0001, unpaired two-tailed t-test). (**O**) Scheme illustrating the experimental setup to induce ablation of Wnt secretion from telocytes within the rostral back skin through the topical application of 4-OH-tamoxifen at the resting phase (p21-p22). (**P-Q**) Representative photographs of mice applied with 4-OH-tamoxifen. Note the fur in the rostral back skin region in a control mouse (P boxed region) compared to the hairless skin in the Foxl1-PorcnΔ mouse (Q boxed region). (**R-V**) Short EdU labeling (white) and immunofluorescence for the SC marker K15 (green) and inner bulge epithelium K6 (red) reveals proliferating follicles and SC expansion in control mice (R, T). In contrast, follicles from Foxl1-PorcnΔ mice show limited proliferation and a restricted bulge compartment (S, U). Quantification of EdU^+^ cells per follicle (V) (n=3 mice per group <20 follicles for each bar ****P<0.0001 unpaired two-tailed t-test). (**W-Y**) Short EdU labeling (white) in intestinal crypts shows no statistically significant difference between control mice compared to Foxl1-PorcnΔ mice (n=3 mice per group <20 crypt for each bar, unpaired two-tailed t-test). Boxed regions highlight the structure of a follicle in each group. Scale bar 100 μm, 25 μm (boxed regions).

Histological analysis revealed that Foxl1-PorcnΔ mice had significantly thinner skin and complete arrest of hair follicle growth compared to controls (**Figures 6C-E**). While control mice exhibited follicles typical of the growth phase, Foxl1- PorcnΔ follicles remained in the resting phase.

Further analysis using EdU labeling combined with P-Cadherin and K15, or Sox9 and K14 staining, revealed that control mice had proliferating hair progenitors and outer root sheath cells, while Foxl1-PorcnΔ mice displayed quiescent follicles arrested in the resting phase (**Figures 6F-K**). Wnt inhibition in telocytes, similar to the effects observed after telocyte ablation, caused a marked reduction in Sox9 expression (**Figures 6I-K**), highlighting the strong dependence of Sox9 on Wnt signaling and its connection to SC activity. Immunohistochemistry for β-catenin showed nuclear localization in progenitor cells at the bulb of control follicles, but in Foxl1-PorcnΔ follicles, β-catenin was restricted to the membrane (**Figures 6L-N**).

To rule out the possibility of intestinal damage, we selectively deleted *Porcn* in skin telocytes by topically applying 4OH-tamoxifen to the back skin (**Figure 6O**). In control mice, hair grew normally in the treated area, while Foxl1-PorcnΔ mice remained hairless in the corresponding region (**Figures 6P-Q**). Follicle proliferation was significantly reduced in Foxl1-PorcnΔ mice, with no observable intestinal effects. Additionally, while control mice showed an expanded K15-expressing SC compartment, Foxl1-PorcnΔ follicles remained in the resting phase (**Figures 6R-Y**).

These results demonstrate that telocytes are a crucial source of Wnt proteins necessary for SC activity during hair follicle regeneration. This role is not compensated by other niche cells lacking Foxl1 expression, highlighting the essential function of telocytes in the hair follicle SC niche.

## Discussion

Homeostasis in regenerative tissues, like the hair follicle and intestine, depends on the precise coordination between SCs and their progeny. Traditionally, the SC niche has been viewed as a localized anatomical domain supporting SC function. However, our study reveals that telocyte networks maintain continuous interaction with both SCs and their progeny throughout follicle regeneration (**Figures 7A-B**). This network consists of dermal telocytes in contact with the basal epithelium and inner telocytes associated with the inner epithelial layers. Some dermal telocytes are part of the dermal papilla in both the resting and growth phases, while during the growth phase, they also contribute to the formation of the dermal sheath. Together, the dermal and inner telocytes form a unified, interconnected network that contacts SCs and their progeny, orchestrating SC activity across multiple follicle compartments. This network undergoes dynamic structural and molecular changes in response to epithelial function, supporting a revised, integrative model of the SC niche, with telocyte networks playing a central role.

**Figure 7.**
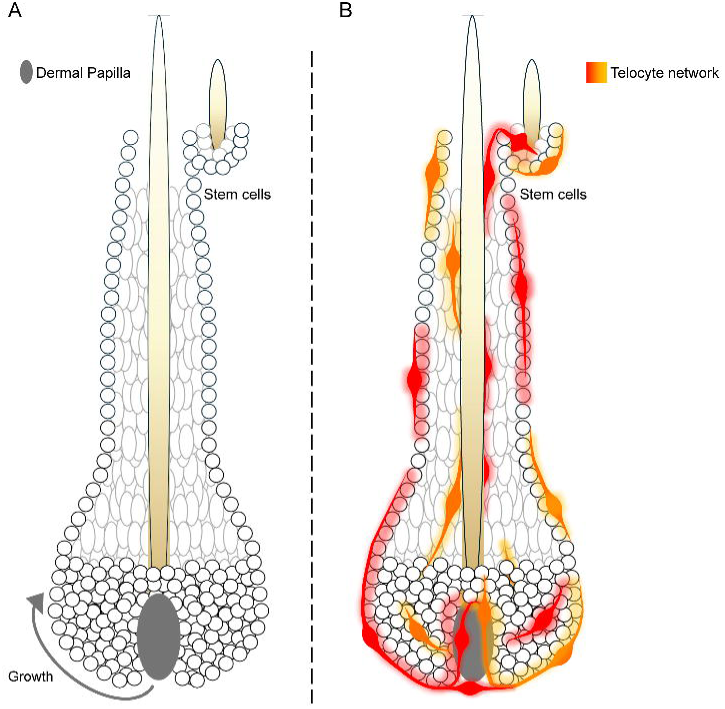
Revised telocyte model of the hair follicle SC niche compared to the dermal papilla model. (**A**) The dermal papilla model proposes that a cluster of mesenchymal cells called the dermal papilla (grey) induces the proliferation of transit amplifying cells located at the follicle bulb, while the epithelium itself coordinates the entire hair cycle. (**B**) In the revised telocyte model, telocyte networks (orange and red) are proposed to continuously coordinate SC activity across follicle layers, providing an integrative, dynamic regulatory framework for hair follicle regeneration.

The hair follicle epithelium is supported by a dense basement membrane that separates it from the dermal compartment. Previously, mesenchymal components of the SC niche were thought to be restricted to the dermis. However, we identified telocytes also in the inner follicle layers, crossing the basement membrane. Remarkably, from the early stages of hair placode morphogenesis, telocytes are present aligning along the developing epidermis and within the nascent hair germ. It has been suggested that the hair follicle develops from concentric 2D zones that evolve into 3D cylindrical layers^110^. Telocytes may extend along the epithelium during this process, establishing themselves in the inner follicle layers. Additionally, dermal telocytes may migrate across the basement membrane as it remodels^111^. Tracking telocytes during placode morphogenesis could provide critical insights into their role in follicle development.

Mesenchymal components involved in follicle development and homeostasis, such as the dermal condensate, dermal papilla and dermal sheath, are typically defined by their location and marker expression. However, the diversity within these components, where telocytes take part, is underappreciated. Ultrastructural studies from the 1960s using electron microscopy described fine fibril networks with beaded patterns around the follicle bulb, dermal papilla and inner root sheath^112,113^. These fibrils were associated with pycnotic, irregular nuclei, similar to those observed in telocytes. Advances in volume electron microscopy now allow for detailed structural characterization of telocyte networks within the follicle.

The continuous subepithelial network of telocytes, which accompanies SCs and their progeny throughout differentiation, suggests a regulatory link between SC activity and progeny differentiation. Electron microscopy has shown that telocyte extensions connect via point contacts and electron-dense nanostructures, with membranes separated by narrow intercellular spaces of 10-30 nm. This suggests the possibility of molecular interactions and the transfer of information along the telocyte network, which may play an important role in regulating homeostasis.

It has been proposed that a subset of dermal sheath cells retained after each hair cycle has self-renewal properties and can repopulate the dermal sheath and dermal papilla during follicle growth^114^. While the telocyte network expands significantly during the growth phase, we did not observe telocyte proliferation in either the resting or growth phases. Since hair follicle growth is tightly regulated by Sonic Hedgehog (Shh) signaling^48,96,115,116^, and Foxl1 is a direct target of this pathway^79^, it is therefore suggesting that telocyte dynamics in follicle regeneration may be regulated by Shh signaling. Further investigation into telocyte expression profiles, morphology, and distribution during the transition from resting to growth phases will clarify the mechanisms driving telocyte dynamics.

Our findings demonstrate that telocytes serve as a SC niche beyond the intestine, acting as a critical source of Wnt proteins essential for SC activity in the hair follicle. This discovery highlights a fundamental mechanism for maintaining homeostasis in regenerative tissues. The challenges in studying these delicate flat cells with their thin membranous extensions, prone to damage during fixation or dissociation, may explain why telocyte networks have been overlooked in other organs. Understanding the role of telocytes in SC function offers a crucial link between individual cell fate decisions and the coordinated maintenance of tissue homeostasis.

## Acknowledgments

We thank all members of M. Shoshkes laboratory for their helpful and insightful discussions. We also thank C. Kalcheim and K.H. Kaestner for their critical reading of the manuscript, and E. Fuchs for her comments and feedback throughout the course of this study. Special thanks to S. Itzkovitz and his laboratory for the assistance with implementing the smFISH technology, and to K.H. Kaestner and Y. Rinkevich for sharing mouse models. Schematic illustrations were created with BioRender.com

This work was supported by the Israel Science Foundation (grant #1997/19, MS-C); Carole and Andrew Harper Diversity Program to Excellent Palestinian PhD Scholarship to A.J; HUJI International PhD Talent Scholarship to A.G;

## Author contributions

M.C wrote the manuscript, conceived, carried out experiments, analyzed and interpreted the data. S.N carried out mouse experiments and performed immunostaining. A. J carried out mouse experiments, N.C and A.G performed smFISH experiments, I.B-P, M.H and M.S.-C designed and supervised the study. M.S.-C wrote the manuscript and directed the study.

## Supplemental information

Document S1.

**Figure S1**. Standard tissue fixation protocols impair the visualization of the Foxl1^+^ cell network.

**Figure S2**. Foxl1 Cre-derived GFP signal faithfully marks Foxl1^+^ cells.

**Video S1**. Foxl1^+^ cells wrap around the inner bulge region at the follicle resting phase.

**Video S2**. At the follicle resting phase, Foxl1^+^ cells are distributed throughout the inter-follicular dermis, beneath the follicle, and along the inner and outer bulge regions.

**Video S3**. The elongated cell bodies of Foxl1^+^ cells are intercalated among the epithelial cuboidal cells.

**Video S4**. Foxl1^+^ cells wrap around the inner bulge region at the follicle full growth phase.

All related to Figure 1.

## STAR Methods

### Mice

#### Foxl1-Cre mice^117^ were crossed with the following lines

Rosa-membrane-targeted dimer tomato protein (mT) or membrane targeted green fluorescent protein (mG) (Rosa-mTmG)^118^ (Jackson Laboratories, Bar Harbor, ME #007676), Rosa-Rainbow^65^ or Rosa-inducible diphtheria toxin receptor mice (iDTR)^119^ (Jackson Laboratories, Bar Harbor, ME #007900). This produced the Foxl1Cre; Rosa- mTmG, Foxl1Cre; Rosa- Rainbow or Foxl1Cre; Rosa-iDTR mouse models.

#### Foxl1-CreERT2 mice^64^ were crossed with

Rosa-mTmG or Porcn-ex3-7Neo-flox^109^ (Jackson Laboratories Bar Harbor, ME #020994). This produced the Foxl1CreERT2; Rosa-mTmG, or Foxl1CreERT2; PorcnΔ mouse models.

All animal experiments were approved by the Animal Care and Use Committee of the Hebrew University of Jerusalem.

### Diphtheria toxin treatment

For Foxl1Cre; Rosa-iDTR mice and Cre-negative Rosa-iDTR controls, diphtheria toxin (Sigma-Aldrich #D0564) dissolved in 0.9% sodium chloride was administered intraperitonially at a dose of 20 ng/g body weight. p20-21 mice received the diphtheria toxin for four consecutive days: once daily for the first two days and twice daily at 8-hour intervals for the next two days.

### Tamoxifen treatment

To induce Foxl1-CreERT2; Rosa-mTmG or Foxl1-CreERT2; PorcnΔ mice, tamoxifen (Sigma-Aldrich #10540-29-1) was dissolved in corn oil at a concentration of 30 mg/ml by shaking overnight at 37°C. The solution was then administered intraperitoneally at a dose of 150 mg/kg body weight for reporter mice or 120 mg/kg body weight for Foxl1-PorcnΔ mice to induce recombination while maintaining intestinal homeostasis. Tamoxifen was given over three consecutive days.

### Topical 4OH-tamoxifen treatment

To induce Foxl1-CreERT2; PorcnΔ mice and CreERT2-negative control mice, 4OH- tamoxifen powder (Sigma #H7904) was dissolved in 100% acetone at a concentration of 7.5 mg/ml. At p20, mice were anesthetized via intraperitoneal injection of 7.5% ketamine / 42.5% xylazine (v/v), and their back skin was shaved. Subsequently, 100µl of the 4OH-tamoxifen solution was applied topically to the rostral back skin of each mouse, once daily for two consecutive days. Mice were then sacrificed and analyzed at p30-33.

### EdU incorporation and labelling

For EdU treatment, mice received an intraperitoneal injection of 100μl EdU (5mg/ml Sigma #900584) per 10 grams of body weight, administered 1.5 hours before they were sacrificed. Detection of EdU was performed using a 10μM Alexa Fluor^TM^647 (Invitrogen # A10277), incubated at room temperature for 30 minutes in a copper-containing buffer (1mM CuSO4, 0.1M TRIS base, 10μM ascorbic acid).

### Skin fixation without perfusion

Mice were euthanized, shaved, and back skin was dissected. The skin was then stretched onto a Whatman paper and fixed in 4% PFA at 4°C overnight.

### Fixation by trans-cardiac perfusion

Mice were anesthetized by intraperitoneal injection of 7.5% ketamine / 42.5% xylazine (v/v) and their back skin was shaved. The thorax and rib cage were then opened to expose the heart. Following this, the right atrium was incised, and a 23G needle connected to a peristaltic pump was inserted into the left ventricle to perfuse pre-cooled (4°C) 4% PFA. Each mouse was perfused with approximately 30 ml fixative. After perfusion, the skin was dissected and further fixed with 4% PFA at 4°C overnight.

### Histology

For hematoxylin and eosin (H&E) staining, paraffin-embedded tissue sections were first rehydrated. Then, slides were incubated with hematoxylin (Sigma #0506002) for 5 minutes, followed by rinsing with water. Subsequently, the sections were briefly immersed in acid ethanol (70% EtOH/ 3% HCl (v/v)) for 10 seconds and washed again with water. Slides were then immersed in 0.2% ammonium hydroxide (Sigma #338810), rinsed once more with water, and immersed in eosin (Sigma #HT110116) for 40 seconds. Finally, the slides were briefly rinsed with tap water, dehydrated and mounted.

### Immunofluorescence and immunohistochemistry

Tissue was fixed by trans-cardiac perfusion, and antigen retrieval was carried out using citrate buffer (pH=6) in a pressure cooker (Electron Microscopy Science). For immunofluorescence, tissue was embedded in optimum cutting temperature compound (OCT), and antibodies were diluted in CAS-Block (Invitrogen 008120).

For β-catenin immunohistochemistry, tissue was embedded in paraffin, followed by antigen retrieval, and antibodies were diluted in Starting block^TM^ (ThermoFisher #37542). Anti-mouse-biotin served as a secondary antibody, VECTASTAIN ABC-HRP Kit, Peroxidase (Vector #PK-4000) was used to detect biotinylated molecules. Finally, 3,3-diaminobenzidine tetrahydrochloride (DAB) was applied as a substrate to develop the signal.

### Antibodies

CD31 (BD Pharmingen #557355 1:250), CD34 (ThermoFisher #14-0341-82 1:100), CD81 (Abcam #104912 1:100), EpCAM (Abcam #71916 1:100), GFP (Abcam #6673 1:400), GFP (NovosBio #100-1614 1:400), K6 (Biolegend #905702 1:400), K14 (Biolegend #906004 1:1000), K15 (Biolegend #833904 1:1000), Mcam (Abcam #34717 1:200), P-cad (R&D systems #AF761 1:100), PDGFRα (R&D systems #AF1062 1:100), Postn (R&D systems #AF2955 1:100), Procr (Biolegend #141505 1:100), aSMA (Abcam #5694 1:250), Sox9 (Milipore #AB5535 1:250), Sox2 (ThermoFisher #14-9811-82 Btjce 1:100), Ngfr (R&D systems #AF1157 1:500), Tnc (Abcam #19011 1:300),b-Catenin (BD Pharmingen #610153 1:200), mouse-Biotin (Jackson Labs #705065147 1:100).

### Clearing of mouse back skin X-CLARITY™

To clear the skin, we utilized Foxl1Cre; Rosa-mTmG mice at p20-21or p30-33. Tissue fixation was performed by trans-cardiac perfusion, followed by clearing the skin for 6 hours using an X-Clarity™ ETC chamber following the manufacturer’s protocol (LOGOS Biosystems, Annandale, VA), as previously described^7^.

### Immunofluorescence of cleared whole tissue

Prior staining, cleared skin was blocked with 10% fetal bovine serum in PBS for 1hour. Primary and secondary antibodies were diluted in a blocking solution containing 0.3% Triton X-100 and incubated for 48 hours at 4°C each.

Finally, the tissue was mounted *en bloc* on an image slide using 0.5mm thick adhesive silicon isolator mounted in Histodenz solution (88% Histodenz (w/v) in 0.02 M PBS).

### Single molecule RNA FISH (smFISH)

We employed Foxl1Cre; Rosa-mTmG mice at p20, p24 or p30. Tissue fixation was performed via trans-cardiac perfusion. Custom probe libraries, designed using the Stellaris FISH probe designer software, were utilized to hybridize with a desired coding RNA (Biosearch Technologies, Inc., Petaluma, CA) and coupled to Cy5.

Cryo-sections of 7-micron thickness were prepared for hybridization according to the protocol^120^ with some adaptations. Briefly, tissue sections were digested with proteinase K (10 µg/ml Ambion AM2546) for 10 minutes and washed twice in 2× SSC (Ambion AM9765). Sections were then washed in wash buffer (20% formamide (Ambion AM9342), 2× SSC) for 5 minutes before hybridization with smFISH probes diluted 1:3000 in hybridization buffer (10% dextran sulfate (Sigma D8906), 20% formamide, 1 mg/ml *E. coli* tRNA (Sigma R1753), 2× SSC, 0.02% BSA (Ambion AM2616), 2 mM vanadyl-ribonucleoside complex (NEB S1402S) at 30°C overnight. Following hybridization, sections were washed with wash buffer containing 100 ng/ml DAPI at 30°C for 30 minutes. GFP antibody was diluted in the hybridization buffer, and Alexa 488 secondary antibody was diluted in the GLOX buffer for 20 minutes, followed by DAPI (Sigma-Aldrich, D9542) staining.

### smFISH quantification

Single molecule mRNA dots were counted along the GFP signal using custom MATLAB program^121^ (MATLAB Release 2024a, The MathWorks, USA). Hair follicles from P20, P24, and P30 mouse back skin was segmented into distinct compartments: isthmus (I), bulge (B), hair germ (HG), dermal papilla (DP), inner bulb (IB) and outer bulb (OB).

### Statistical analysis

Statistical and graphical data analyses were conducted using GraphPad Prism 9.5.1. The number of replicates for each experiment and the statistical tests performed are detailed in figure legends.

### Microscopy

Microscopes used in the study:

- Yokogawa w1 spinning disk attached to an inverted Nikon Ti2E microscope (Nikon, Japan).
- Nikon Eclipse Ti2 Confocal System (Nikon, Japan).
- Nikon TI2-LA-FL EPI-FL module for smFISH technology (Nikon, Japan).
- Nikon DS-Fi2 color CCD camera attached to a Nikon Ti automated inverted microscope (Nikon, Japan).

Image processing was performed using Fiji, Imaris v10.0 or NIS Elements software packages.

### Software

Seurat v5.1.0^122^ and DESeq2^123^ were run on R version 4.3.2.

### Data resources and data analysis

The datasets analyzed in this study include GSE122026 (fetal mouse E15.0 bulk and single-cell RNA-sequencing)^76^, the Tabula Sapiens sequencing data of human skin^98^ and the Cellxgene sequencing data of human stromal cells^99^. Bulk RNA sequencing data was analyzed in an R Environment (RStudio 2024.04.2-764) using DESeq2 (Bioconductor 3.19). Single-cell RNA sequencing datasets were analyzed using Seurat v5, with the default parameters and the recommended analysis protocols, in an R Environment (R 4.4.0, RStudio 2024.04.01 +748).

