## Supplementary material for "Telocyte networks are an essential stem cell niche component in hair follicle regeneration": Document S1.docx

Supplemental information

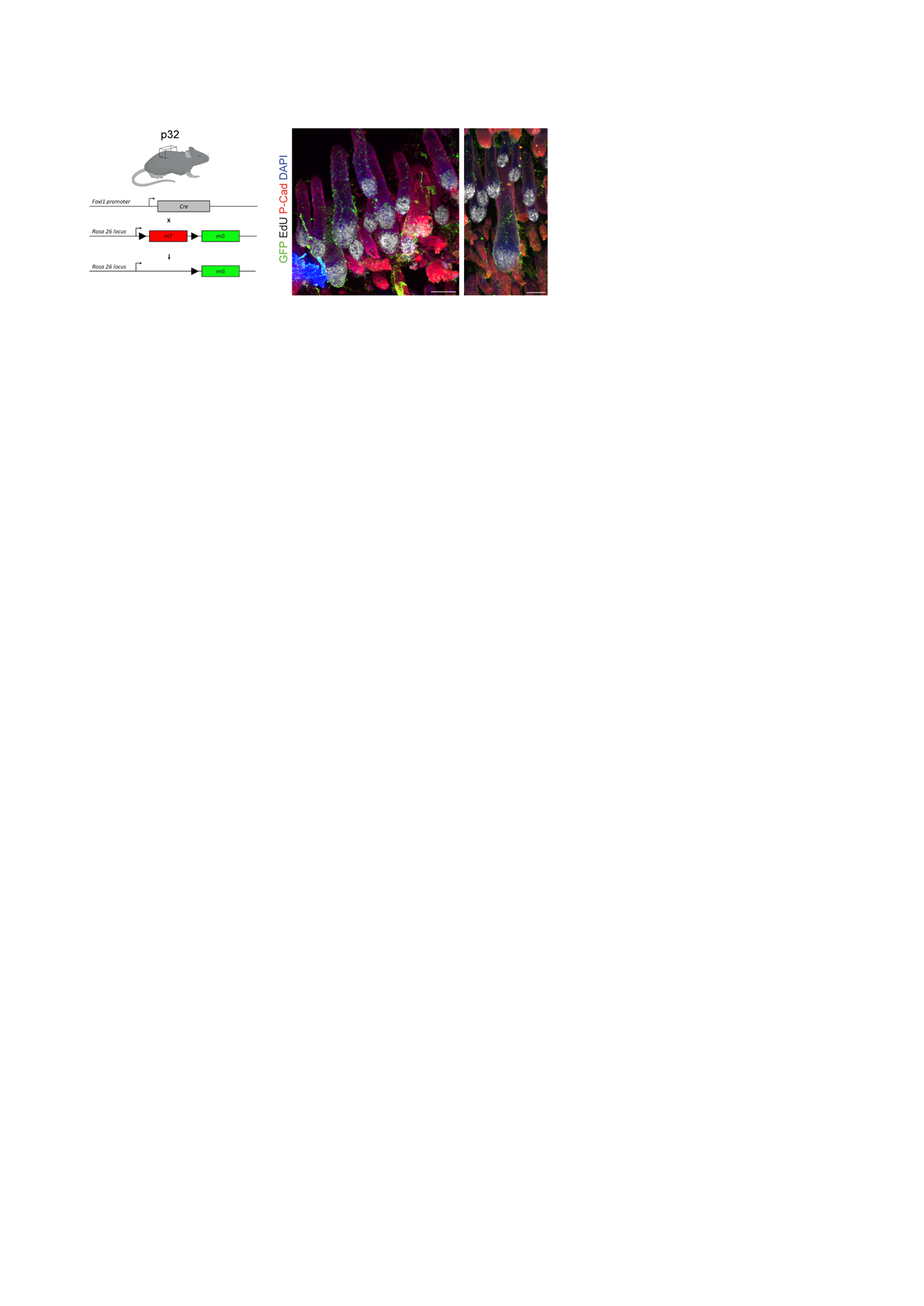

Figure S1. Standard tissue fixation protocols impair the visualization of the Foxl1^+^ cell network.

Whole-mount immunofluorescence of p32 Foxl1-Cre; Rosa-mTmG mouse back skin, following standard fixation protocol and tissue clearing, shows a dramatic impairment in the visualization of the GFP signal. EdU incorporation (white), P-Cad (red), Scale bar 100μm.

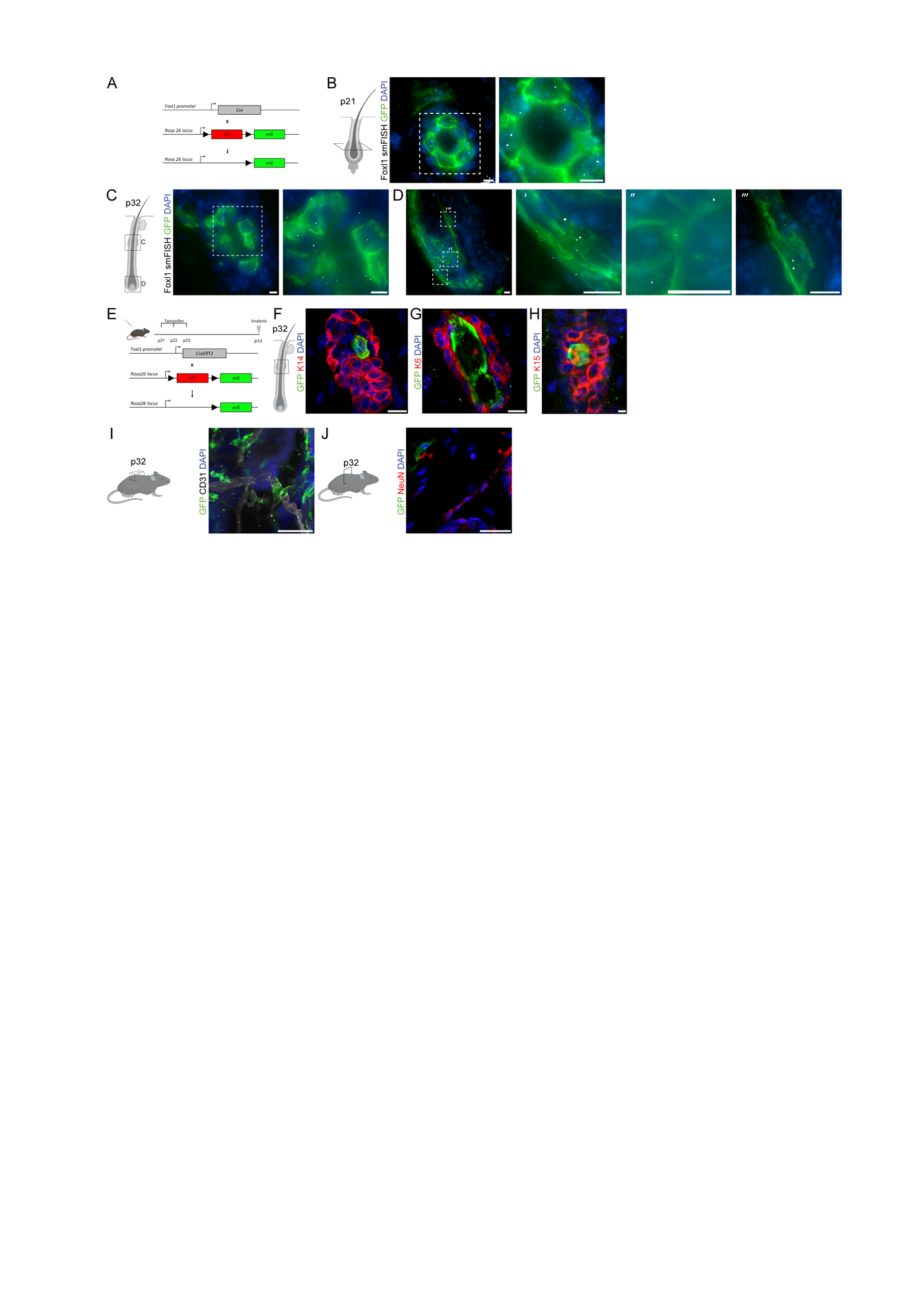

**Figure S2. Foxl1 Cre-derived GFP signal faithfully marks Foxl1^+^ cells**

(**A**) Scheme of mouse model illustrating how the expression of membrane-bound GFP is driven in Foxl1-Cre-positive cells in the Foxl1-Cre; Rosa-mTmG mouse model. (**B-D**) smFISH images show cross and longitudinal sections of the p21(B) and P32 (C-D) bulge (B-C) and bulb (D) regions, hybridized to detect Foxl1 mRNA molecules (grey dots). Immunofluorescence for GFP reveals the expression of Foxl1 mRNA transcripts in GFP-labelled cells. Boxed regions highlight Foxl1^+^ cells in the bulge and bulb regions. Scale bar 5μm. (**E-H**) Scheme illustrating the experimental setup in which tamoxifen induction drives the expression of membrane-bound GFP in Foxl1-Cre-positive cells in the Foxl1-CreERT2; Rosa-mTmG mouse model (E). Mice were induced at p21 and were analyzed at p32. Immunofluorescence of p32 follicles reveals the induction of GFP expression in cells along the inner bulge region stained for K14 (F), K6 (G), and K15 (H). Scale bar 10μm. (**I-J**) Immunofluorescence of skin whole mount and section from p32 Foxl1-Cre; Rosa-mTmG mouse model reveals Foxl1^+^ cells closely associated with blood vessels, stained for Cd31(I), and neurons stained for Neun (J). Scale bar 50μm.

Video S1.

**Foxl1**^+^ **cells wrap around the inner bulge region at the follicle resting phase.**

Immunofluorescence of a 20μm longitudinal section of p21 Foxl1Cre; Rosa-mTmG mouse back skin reveals the distribution of Foxl1^+^ cells along the inner bulge region, where stem cells stained for K15 (red) reside, during the follicle resting phase.

**Video S2.**

**At the follicle resting phase, Foxl1**^+^ **cells are distributed throughout the inter-follicular dermis, beneath the follicle, and along the inner and outer bulge regions.**

Immunofluorescence of a 20μm longitudinal section of p21 Foxl1Cre; Rosa-mTmG mouse back skin reveals the distribution of Foxl1^+^ cells throughout the inter-follicular dermis, beneath the follicle and along the outer and inner bulge region where stem cells stained for K15 (red) reside.

**Video S3.**

**The elongated cell bodies of Foxl1**^+^ **cells are intercalated among the epithelial cuboidal cells.**

Immunofluorescence of a 20μm longitudinal section of p24 Foxl1Cre; Rosa-mTmG mouse back skin reveals that the Foxl1 DAPI+ nuclei exhibit an elongated structure with puncta of heterochromatin regions, distinct from the cuboidal epithelial nuclei.

**Video S4.**

**Foxl1**^+^ **cells wrap around the inner bulge region at the follicle full growth phase.**

Immunofluorescence of a 20 μm cross section of p30 Foxl1Cre; Rosa-mTmG mouse back skin reveals that the Foxl1^+^ cells envelope the inner bulge region, where K15 stem cells, stained in red, reside during the growth phase.
